# Gut MicrobiotAware: how much do we know about gut microbiota? An international questionnaire

**DOI:** 10.1101/2024.12.04.626811

**Authors:** Enriqueta Garcia-Gutierrez, Sara Arbulu, Charlotte Oliver, Sandeep Kumar, Sarita A. Dam, Babette Jakobi, Vincenzo Pennone, Fabiana A. Hoffmann Sarda, Arghya Mukherjee, Paul D. Cotter

## Abstract

Over the past two decades, our understanding of human gut microbiota composition and function has grown significantly. The surge in studies and data has unveiled the profound influence of gut microbiota on host nutrition, health, and behavior, bridging biology, medicine, and ecology. The dynamic interaction between daily choices, life events, and gut microbiota composition makes it a fascinating subject for scientists to communicate to diverse audiences. Effective scientific communication involves adapting the delivery of knowledge to different audiences, using precise language in academic settings and accessible concepts in public forums like science festivals or social media. With the growing interest in gut microbiota beyond academic circles, there is an increased risk of disseminating information lacking scientific rigor. This study aims to assess the general public’s knowledge about gut microbiota and evaluate healthcare professionals’ understanding of its links to various health conditions, ultimately informing better communication strategies for both groups.

## 1. Introduction

The contribution to the health of the human gut microbiota, i.e., the collection of microbes that live in our gastrointestinal tract, includes bacteria (the most studied group), archaea, viruses, fungi and protozoa, has become increasingly apparent over the last two decades [1,2]. Disruption of the gut microbiota or specific gut microbial signatures has been linked to many gastrointestinal conditions, obesity, type 2 diabetes, neurological disorders such as Parkinson’s and Alzheimer’s, asthma, rheumatoid arthritis and even cancer [3–10]. As a result, the gut microbiome has emerged as a therapeutic target, prompting efforts to develop tools to understand and modulate this ecosystem in ways that can translate into patient care.

Discussions relating to the gut microbiota have moved beyond exclusively academic circles to the general public, permeating different demographic groups, and the healthcare system. The field has been the focus of considerable efforts relating to science dissemination and communication [11,12].

Science communication refers to the processes and practices used to share scientific knowledge with the general public or non-expert audiences [13]. On the other hand, science dissemination refers to making scientific findings available to expert audiences, including the scientific community and professionals in the field, the industry and policymakers. Depending on which group is being addressed, different approaches, activities and language use are required. While disseminating within the scientific community implies using precise, unambiguous language (e.g., scientific articles and conferences), conveying the same information to the general public requires accessible concepts and language (e.g., science festivals, podcasts or social media). When relaying information relating to the gut microbiota, it is essential to transmit this knowledge rigorously to raise awareness and educate not only the public but also health professionals, as they are the crucial players in our health systems.

Current efforts to develop tools for assessing gut microbiota as part of overall health are still in progress, and routine analysis and interpretation of the gut microbiota at the patient level are not yet fully available to the general population [14]. This is significant because the gut microbiota’s composition varies with factors such as diet, lifestyle, and geographical location, making it difficult to establish a universal healthy gut microbiota standard [15–17]. Furthermore, while there are various services claiming to analyse the gut microbiota, the quality and the accuracy of their claims vary, reflecting ongoing debates in the field. Moreover, to fully harness the potential of the gut microbiota, it is necessary to address knowledge gaps, particularly in understanding interactions between microbes and between microbes and their host. This includes gaining a deeper understanding of less studied organisms such as fungi and viruses, getting insights into microbial functions in addition to descriptive composition, conducting large-scale studies and using all this information to understand this complex ecosystem [17].

This work aimed to assess the general public’s and healthcare professionals’ knowledge relating to gut microbiota concepts. The survey covered questions regarding the level of general interest among the responders, the sources of media used to learn about the gut microbiota as well as the specific knowledge of the participants concerning technical concepts. Additionally, we queried to what extent gut microbiota analyses are in routine practice among healthcare professionals. Ultimately, this information will help to design more effective strategies to communicate gut microbiota research.

## 2. Methods

### 2.1. Questionnaire design

This cross-sectional study was conducted between April 2022 and November 2023, using an online questionnaire via the SurveyMonkey platform. The questionnaire consisted of 22 or 23 questions (for the United States of America, India and China, an additional question regarding province location was included) grouped in three blocks: i) demographic information; ii) general knowledge relating to the gut microbiota and its connections with human health and iii) application of gut microbiota-related knowledge in the healthcare sector. The questionnaire was designed in English and translated into Bengali, Chinese, Dutch, French, German, Hindi, Italian, Portuguese, Punjabi and, Spanish by native speakers who are also proficient in English. Answers were selected from multiple choice options and opt-outs like “I do not know”, “I am not X”, “I prefer not to respond” were provided. Research suggests that including such options improves completion levels [18,19].

### 2.2. Participant recruitment

Participation was voluntary, independent and without compensation. Neither personal information nor IP addresses were collected, making the responses completely anonymous. The questionnaire was distributed via different channels: i) university communities and networks of collaborators; ii) social media and email lists of research funding bodies and other organisations; iii) social media groups and iv) family and friends networks.

### 2.3. Statistical analysis

Statistical analyses were conducted using GraphPad Prism 8 and R version 4.4.1 (2024-06-14). Answers were manually curated to identify potential inconsistencies within questionnaires (e.g., answering to healthcare answers if previously identified as non-healthcare workers). The final number of curated responses is indicated in each figure.

## 3. Results

### 3.1. Socio-demographics

A total of 1289 participants from 54 countries across the five continents completed the survey between March 2022 and November 2023. Spain was the country with the largest number of participants (426), followed by Ireland (149), China (144) and Brazil (98) (Table 1). A total of 41 countries had less than 10 participants.

**Table 1.** Number of participants by country.

| Table 1. Number of participants by country. |  |
| --- | --- |
| Country of origin | Number of participants |
| Algeria | 7 |
| Andorra | 1 |
| Anguilla | 1 |
| Argentina | 7 |
| Australia | 8 |
| Austria | 4 |
| Bangladesh | 7 |
| Belgium | 8 |
| Bolivia | 1 |
| Brazil | 98 |
| Bulgaria | 1 |
| Canada | 16 |
| Chad | 1 |
| Chile | 7 |
| China | 144 |
| Colombia | 6 |
| Czech Republic | 2 |
| Denmark | 4 |
| Ecuador | 3 |
| Ethiopia | 1 |
| Finland | 4 |
| France | 29 |
| Germany | 28 |
| Ghana | 2 |
| Guatemala | 2 |
| India | 49 |
| Ireland | 149 |
| Israel | 2 |
| Italy | 35 |
| Malaysia | 2 |
| Mexico | 1 |
| Netherlands | 47 |
| New Zealand | 4 |
| Nigeria | 4 |
| Norway | 7 |
| Pakistan | 1 |
| Palau | 1 |
| Peru | 4 |
| Poland | 3 |
| Portugal | 4 |
| Romania | 1 |
| Saudi Arabia | 2 |
| Singapore | 1 |
| Slovenia | 1 |
| South Africa | 2 |
| Spain | 426 |
| Sweden | 2 |
| Switzerland | 20 |
| Serbia | 1 |
| Turkey | 2 |
| United Kingdom | 60 |
| United States | 55 |
| Venezuela | 1 |
| Zambia | 1 |

Participants’ ages ranged from 18 to over 50 years old with different educational backgrounds and professions grouped as health- or non-health-related (Table 2). Students were assigned to the job category that they were conducting their studies on.

**Table 2.** Socio-demographic characteristics of the participants.

| Table 2. Socio-demographic characteristics of the participants. |  |  |  |
| --- | --- | --- | --- |
|  | Options | Number of responses | Percentage (%) |
| **Age range** |  |  |  |
|  | 18-20 | 28 | 2.19 |
|  | 21-29 | 361 | 28.22 |
|  | 30-39 | 357 | 27.91 |
|  | 40-49 | 253 | 19.78 |
|  | 50-59 | 183 | 14.31 |
|  | 60 or older | 98 | 7.66 |
| **Highest level of education completed** |  |  |  |
|  | Primary education | 7 | 0.55 |
|  | High school | 103 | 8.13 |
|  | Bachelor's degree | 344 | 27.15 |
|  | Master's degree | 373 | 29.44 |
|  | PhD, MD or similar | 372 | 29.36 |
|  | Other | 87 | 6.87 |
|  | Prefer not to say | 13 | 1.03 |
| **Healthcare professional or scientist** |  |  |  |
|  | Medical doctor General Practitioner | 3 | 0.33 |
|  | Medical doctor Gastroenterologist | 2 | 0.22 |
|  | Medical doctor Neurologist | 4 | 0.44 |
|  | Medical doctor specialist (other categories) | 34 | 3.74 |
|  | Nurse | 28 | 3.07 |
|  | Nutritionist | 51 | 5.6 |
|  | Microbiologist | 141 | 15.5 |
|  | Food technologist | 54 | 5.93 |
|  | Pharmacist | 31 | 3.4 |
|  | Veterinarian | 14 | 1.53 |
|  | Other researcher/scientist | 157 | 17.25 |
|  | Retired from any of the mentioned categories | 6 | 0.66 |
|  | I was a healthcare professional or researcher but switched careers | 18 | 1.32 |
|  | I prefer not to say | 12 | 13.74 |
|  | Other | 445 | 39 |
| **Not a healthcare or scientist** |  |  |  |
|  | Architecture and engineering occupations | 42 | 5.49 |
|  | Arts, design, entertainment, sports, and media occupations | 30 | 3.92 |
|  | Business and financial occupations | 35 | 4.57 |
|  | Cleaning and maintenance occupations | 8 | 1.04 |
|  | Community and social service occupations | 21 | 2.74 |
|  | Computer and mathematical occupations | 64 | 8.37 |
|  | Construction and extraction occupations | 2 | 0.26 |
|  | Education, training and library occupations | 107 | 13.99 |
|  | Farming, fishing and forestry occupations | 62 | 8.10 |
|  | Food preparation and serving related occupations | 52 | 6.8 |
|  | Installation, maintenance and repair occupations | 3 | 0.39 |
|  | Legal occupations | 9 | 1.17 |
|  | Management occupations | 33 | 4.31 |
|  | Office and administrative support occupations | 61 | 7.97 |
|  | Personal care and service occupations | 3 | 0.39 |
|  | Production occupations | 9 | 1.17 |
|  | Protective service occupations | 2 | 0.26 |
|  | Sales and related occupations | 23 | 3 |
|  | Social science occupations | 14 | 1.83 |
|  | Transportation and materials moving occupations | 5 | 0.65 |
|  | Other | 133 | 17.38 |
|  | Prefer not to say | 47 | 6.14 |

In the health-related group, ‘Microbiology’ was the most reported occupation, along with researchers and scientists working in other categories with non-provided options. In the non-health group, ‘Food preparation’, ‘Farming, fishing and forestry’ and ‘Education, training and library’ occupations were the most reported options across all ages.

### 3.2. Sources of information and preferred learning channels on gut microbiota

A large number of participants (85.64%) had heard about the gut microbiota, 4.21% had not, and 10.15% were not sure (Figure 1a). The proportion of health-related (45.66%) and non-health-related respondents (54.34%) who replied “Yes” to the question was similar. However, the proportion of non-health-related respondents (92.68%) was higher among the participants that replied that were unsure about hearing the term gut microbiota (Figure 1a).

**Figure 1.**
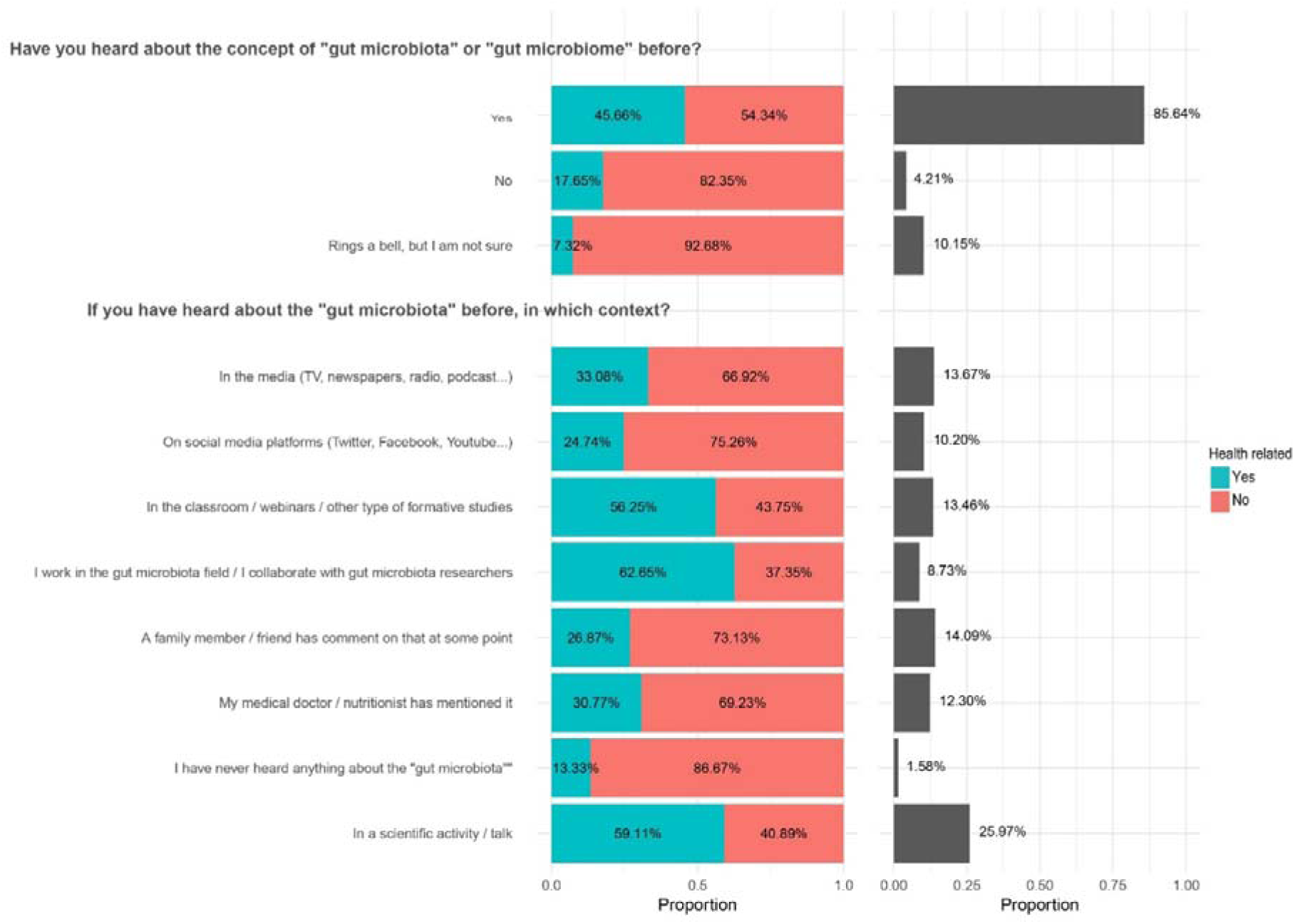
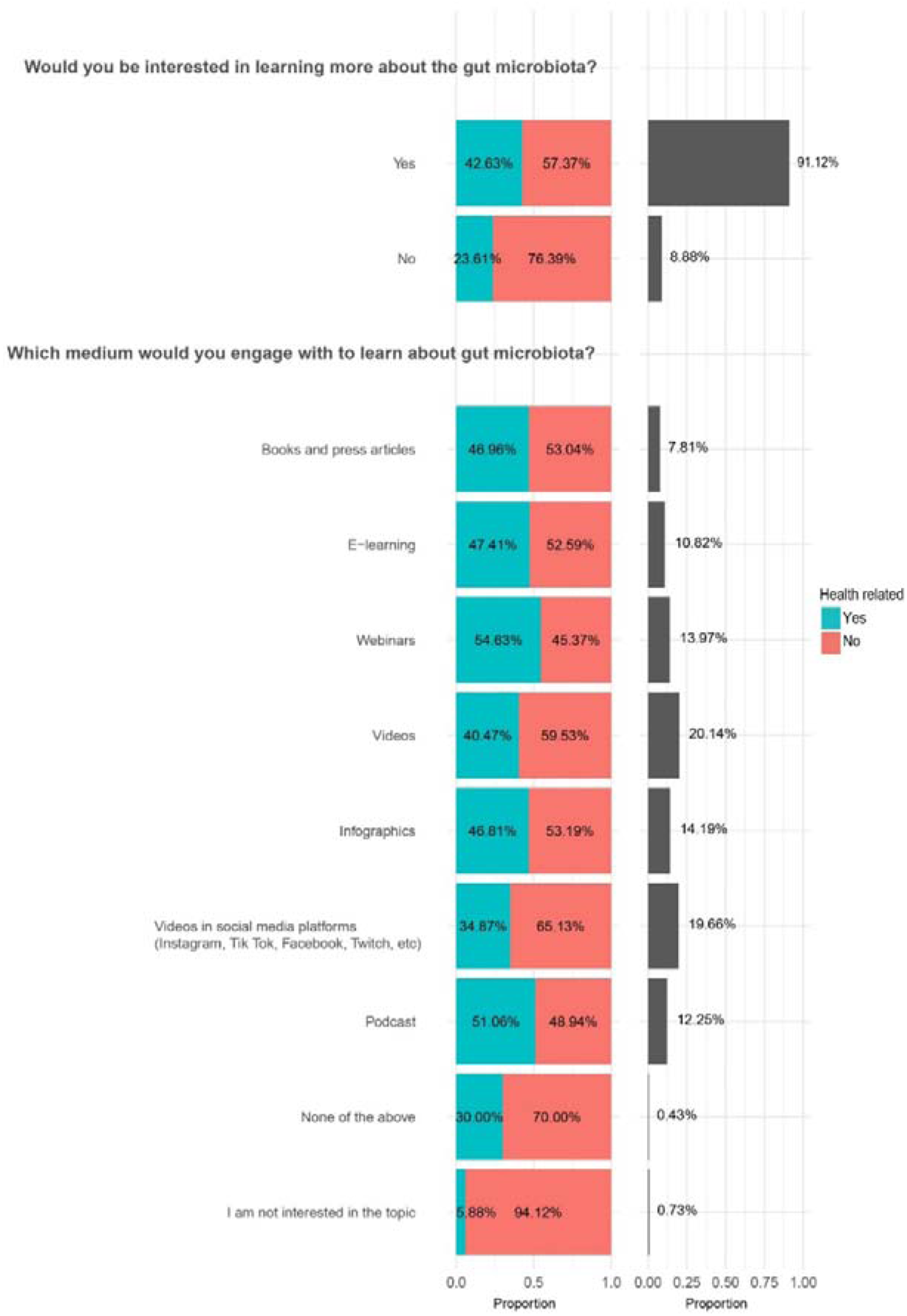
Sources of information and preferred learning channels on gut microbiota. (1a) Sources of information used by the participants (number of responses for each question in order of appearance: n = 808, n = 555). (1b) Sources of information the participants would be willing to use (number of responses for each question in order of appearance n = 811, n = 791).

Where relevant, when the participants were asked where they had heard about the gut microbiota concept, scientific activities/talks was the most selected answer (25.97%) being chosen in higher proportion by health-related respondents (59.11%). All the other options, including social media, health advisors, training and education or word of mouth with family and friends ranged from 10.20% to 14.09% of the answers, indicating the potential importance of using multiple different routes to disseminate information (Figure 1a).

Figure 1b shows the most commonly used media through which individuals learned about the gut microbiota. Videos (20.12%) and videos on social media platforms (19.69%) were the most selected options, followed by infographics (14.19%), while books and press articles where the least popular selection (7.79%). Webinars (13.98%), podcasts (12.25%) and e-learning (10.82%) were other alternatives that respondents would choose to learn about the gut microbiota. Both health and non-health-related participants shared similar rates of affinity for the different options (Figure 1b).

### 3.3. General knowledge of gut microbiota concepts and connection to human health

Six questions pertaining to terms and concepts relating to gut microbiota and health were included in the survey (Figure 2). Participants associated “Bacteria” with gut microbiota in 29.25% of the cases. The second concept that was most associated with gut microbiota was “Faecal matter” (18%) and the third was “Fungi” (10.85%), closely followed by “Mucus” (9.68%), “Epithelium” (9%) and “Viruses” (8.56%). “Protozoa” (6.72%) and “Archaea” (6.08%) were less associated with gut microbiota. Health and non-health related respondents answered similarly to this question. A small percentage of respondents reported not knowing (1.51%) or did not associate any of these terms to the gut microbiota (0.29%). Most of those were from non-health-related occupations.

**Figure 2.**
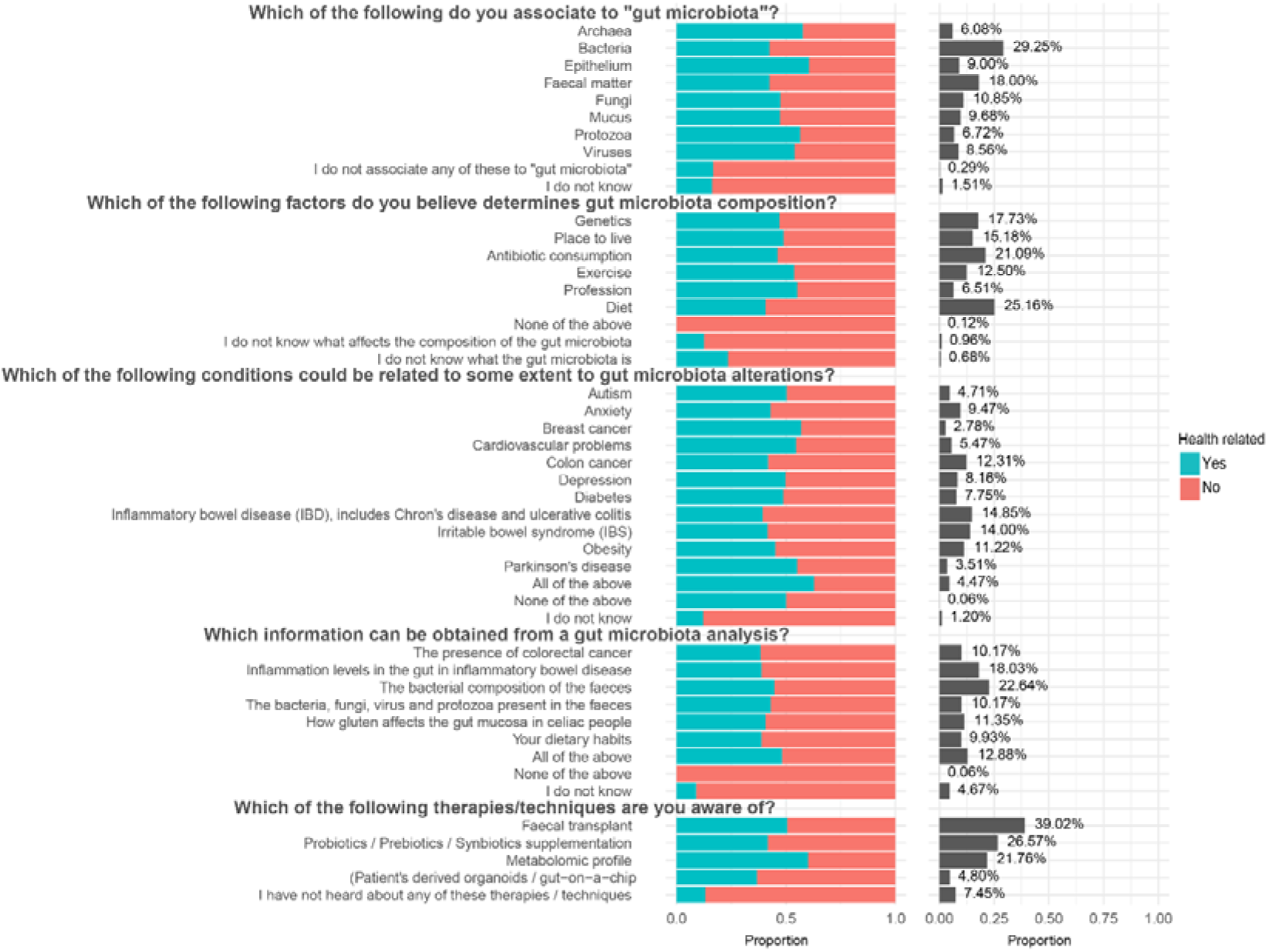
General knowledge of gut microbiota-related concepts (number of responses for each question in order of appearance n = 674, n = 674, n = 672, n = 653, n = 592).

Among the factors that affect gut microbiota composition, diet (25.16%) and antibiotic consumption (21.09%) were the most reported ones, while others like genetics (17.73%), place to live (15.18%), exercise (12.50%) and profession (6.51%) were less associated to the gut microbiota. As for the previous question, similar responses were obtained from both health and non-health-related respondents. In the opt-out options, a small percentage of primarily non-health respondents reported not knowing what affects the composition of the gut microbiota (0.96%) or not knowing what the gut microbiota is (0.68%). Only 0.12% (non-health-related) selected that none of the factors affected the gut microbiota.

When participants were asked which conditions they believed were associated with the gut microbiota, gut conditions such as inflammatory bowel disease (IBD), including Crohńs disease and ulcerative colitis (14.85%), irritable bowel syndrome (IBS) (14%) and colon cancer (12.31%) were among those most frequently selected, while conditions that were not directly connected with the gut, like cardiovascular problems (5.47%), Parkinsońs disease (3.51%) and breast cancer (2.78%) were less associated with the gut microbiota. Systemic conditions like obesity (11.22%) and diabetes (7.75%) were selected to a similar degree as conditions associated with the brain/nervous system such as anxiety (9.47%) and depression (8.16%), whereas autism (4.71%) was not as perceived to be as highly associated with the gut microbiota. 4.47% believed that all conditions were associated to a certain extent with the gut microbiota and only 0.06% did not believe that the gut microbiota had a role in any of the conditions listed. Following the pattern for previous questions, both health-related and non-health-related respondents provided comparable patterns of responses. However, non-health-related respondents more frequently selected “I do not know” (4.67%).

We were also interested in learning what participants perceived they would be provided with through a gut microbiota analysis. The most selected option was the bacterial composition of the faeces (22.64%), although other options that were frequently selected were: inflammation levels in the gut in inflammatory bowel disease (18.03%), how gluten affects the gut mucosa in celiac people (11.35%), inform of the presence of colorectal cancer (10.17%) and dietary habits (9.93%). Moreover, 12.88% of responders believed such an analysis could provide information of relevance across all of these areas, while only 0.06% of non-health respondents answered it couldn’t provide any of these insights. As observed before, there were similar percentages of health and non-health respondents when selecting the options, but we observed the same trend of the latter group being more likely to select the opt-out answer “I do not know” option (4.67%).

Regarding how to order a gut microbiota analysis, visiting a gastroenterologist or ordering it from a private company were selected in 42.69% and 28.22% of cases, respectively, while the family doctor got 16.52% of the answers. Not knowing how to order a gut microbiota analysis was selected in 12.13% of cases.

The last question of this block referred to therapeutic options that are currently under research or have recently started to be available for patient care. The answers of the respondents were as follows: faecal transplants (39.02%), probiotics, prebiotics and synbiotic supplementations (26.57%), metabolomic profiling (21.76%) and organoids and gut-on-a-chip technology (4.80%). 7.45% of responders were not aware of these innovations.

Overall participants answering “None of the above” or “I don’t know” were non-health related participants.

### 3.4. Interest among healthcare professionals in gut microbiota analysis

This block of four questions was designed to assess if microbiota awareness is translated into medical practice (Figure 3). There is a percentage of health professionals that very frequently (26.62%) or often (34.42%) receive patients who suffer from persistent abdominal pain and feeding problems that do not improve with dietary changes. A total of 38.96% of the healthcare professionals recommended a gut microbiota analysis, while 61.04% did not. Those that recommended a gut microbiota analysis did it in the same centre where the healthcare worker was based or a subsidiary one (19.51%) or as part of a clinical study (17.68%). A 16.46% did it via a research centre they collaborated with, while 13.41% used a private company. The health professionals who did not recommend a gut microbiota analysis referred to not having the facilities or resources as the main reason for not recommending one (28.90%), while 16.18% thought that the obtained information would not be relevant. Not considering it (13.29%) and pricing (13.87%) were the next most selected reasons. Finally, other reasons included technical factors such as preferring to conduct different types of analyses that could be more informative or relevant for the patient’s condition (9.25%), believing that the gut microbiota was not involved in the clinical profiles that they were working with (2.31%) or not knowing how to interpret the results of gut microbiota analyses (8.67%).

**Figure 3.**
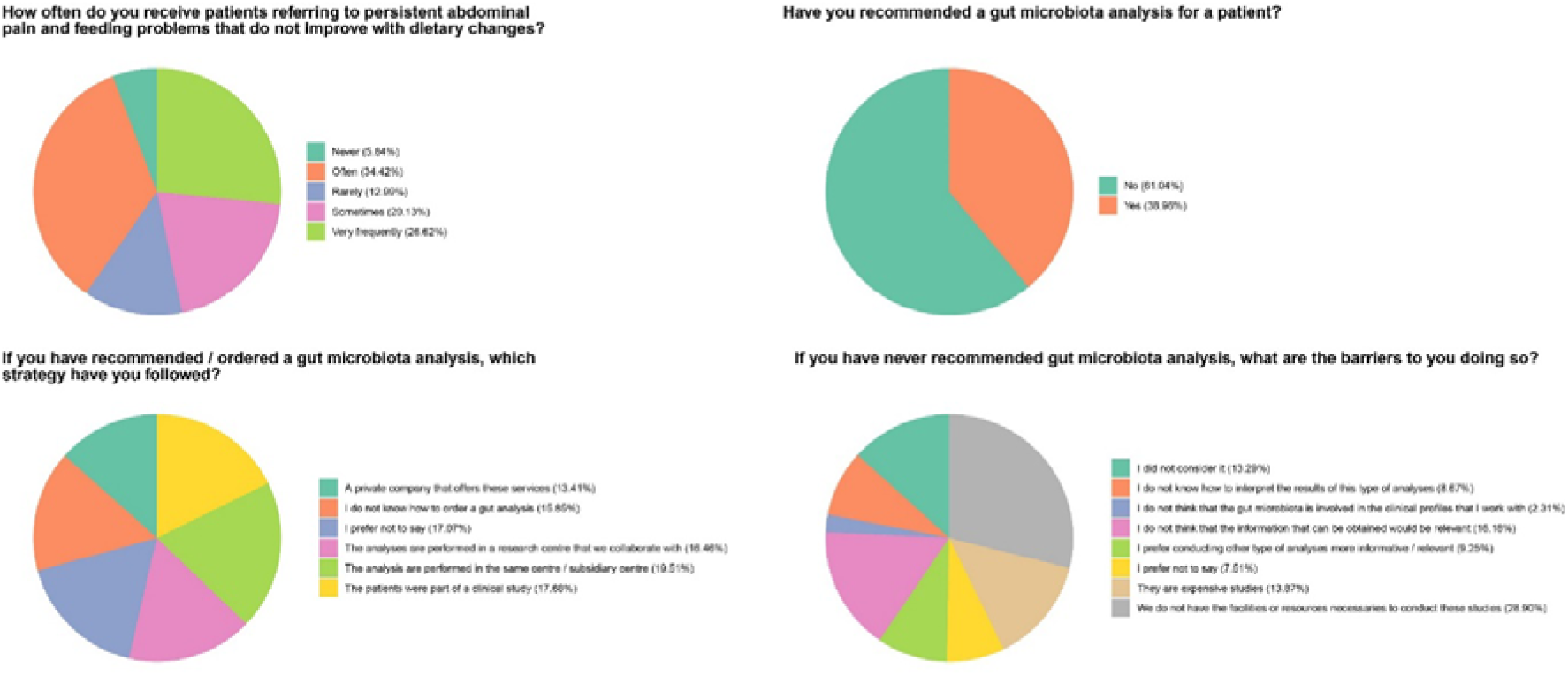
Healthcare professionals’ tendency towards gut microbiota analysis (number of responses for each question in order of appearance n = 154, n = 154 n = 95 n = 107).

### 3.5. Additional questions

In addition to the main results described above the supplementary material includes the results of four additional questions. The participants were asked whether the gut microbiota affects health and mood, and a majority of the participants answered yes in both cases (93.97% and 82.86%, respectively) (Figure S1). When asked where they searched for gut microbiota information research articles (46.61%) and social media (16.25%) were among the most searched channels (Figure S2).

## 4. Discussion

This study reports the results of an online microbiota awareness survey conducted over 19 months between 2022 and 2023 aimed at the general public and health professionals. A total of 1289 participants from 54 countries completed the questionnaire. The sampled population was higher in some countries, such as Spain, China and Brazil, and 41 countries had less than 10 representatives. This does not allow a deep study of the dissemination strategies in the different countries but provides an overview of how widespread certain microbiota concepts are. It is noted that there is an overrepresentation of the number of PhD, MD or similar-educated participants (29.36 %). These numbers are influenced by the survey circulation channels that included universities and research centres, which resulted in a large percentage of participants having undertaken higher education. In a global context, PhD holders only account for 1 % of the world population [20], which sets the context for the education range. Note that health-related jobs included clinical and research occupations that could be, but not necessarily, linked to the gut microbiota. For example, microbiologists are research scientists but not all of them work within the gut microbiota field and, therefore, might be less familiar with the topic.

Given that healthcare and non-healthcare related professionals participated in the questionnaire, the survey gave the opportunity to compare the extent to which gut microbiota-related knowledge is spread among the general population relative to health-care professionals, the latter being increasingly important players for applying gut microbiota knowledge. The obtained results allowed us to identify knowledge gaps and misperceptions about the gut microbiota.

The gut microbiota research field has become a rapidly evolving area of research in medical sciences during the last two decades, with important implications for human health [21]. This new knowledge transcends the boundaries of the research community in the field to reach society at all levels. Unsurprisingly, almost 90% of the respondents had heard about the gut microbiota concept through numerous channels including scientific talks, family members/friends, media, formative studies and family doctors and nutritionists. This is in agreement with The International Microbiota Observatory survey 2024 that reported 70% of the participants had heard about the microbiome term mainly via healthcare professionals, schools, and TV shows [22] (note that the terms microbiota (collection of microbes) and microbiome (collection of microbes and their genes) are used interchangeably in some of these studies [23]). Participants learnt about gut microbiota and wanted to continue learning using a variety of information channels highlighting their interest in the topic and multiple options to deliver effective gut microbiota research outreach, such as videos, social media platforms, infographics, webinars or podcasts.

Knowledge about the composition, factors influencing the gut microbiota, and related-conditions were mainly associated with concepts or conditions directly related to the gastrointestinal tract. This was also observed in a gut microbiota survey targeted at dietitians [24]. In terms of composition of the gut microbiota, the answers were biased towards bacteria (29.25% of the answers). This is not surprising since the bacteriome is by far the best characterised component of the gut microbiota [25]. Viruses, fungi, archaea and eukaryotic organisms are now slowly gaining more attention [26,27]. The term “Faecal matter” was associated with the gut microbiota in 18% of cases. Most of the gut microbiota composition analysis are performed using faecal samples due to their easy collection and processing. Importantly, while faecal matter serves as an indicator of gut microbiota composition, it is not representative of all sub-habitats along the gastrointestinal tract, access to which requires more invasive and costly biopsy-like procedures [28] or newer gut content-capturing capsule technologies [29,30].

Interestingly, diet and antibiotics were regarded as the two main factors that influence the gut microbiota, which is consistent with a previous survey in which 76% and 67% of the participants also selected these two factors, respectively, as being of key importance (BIOCODEX Microbiota Institute, 2024). Research has shown that low microbiota diversity is associated with inflammatory diseases and metabolic disorders [31] and eating a wide variety of foods, especially fermented foods and fibre-rich and exercise lifestyle supports a healthy gut microbiota [32–36]. In addition, numerous campaigns have drawn attention to the antimicrobial resistance crisis and the appropriate use of antibiotics [37,38]. Antibiotics can directly alter the gut microbiota composition decreasing its diversity [39]. Other factors such as genetics, place to live, exercise or profession were selected, highlighting a good knowledge of the variety of factors that can influence the gut microbiota among the participants.

The conditions regarded as being related to the gut microbiota were those associated with gastrointestinal health, e.g., IBD, IBS, colon cancer or obesity. Awareness of other areas where an increasing amount of research is being performed, such as those relating to the gut-brain axis, could benefit from continued science communication.

The use of gut microbiota analysis is not widespread in clinical practice, and this was reflected in the participants’ answers. Although most respondents identified doctors or private companies offering gut microbiota tests, 12% did not know how to order a test. Their view was that such tests can give information on various gut-related conditions.

When healthcare workers were asked about gut microbiota analyses in their routine practice, the fact that 53.65% of the means conducted to perform the gut microbiota analyses derived from analyses in the same centre /subsidiary centre where they were based (19.51%), patients already being part of a clinical study (17.68%) or performed by a collaborator centre (16.46%), suggests that these gut microbiota studies are conducted on the basis of opportunity, i.e., due to availability of resources and/or collaborators with resources. Moreover, from the 61.04% of the healthcare workers who reported that they had never recommended a gut microbiota analysis, 51.44% identified some level of resource limitation, facilities (28.90%), either economic (13.87%) or knowledge-related (8.67%). It is indicative that a 38.96% have not recommended gut microbiota analyses or have not considered it (13.29%), highlighting a potential area of intervention with informative purposes. The debate around gut microbiota testing involves several key issues, reflecting both the promise and limitations of this emerging field [40]. Accuracy and reliability of gut microbiota tests, especially at-home-kits, can lead to varying results from the same stool sample, suggesting that current testing technologies may not provide an accurate snapshot of an individual’s microbiome health [41]. Additionally, being the gut microbiota highly dynamic and influenced by diet, sleep, and stress [42,43], rapid changes can result in tests becoming quickly irrelevant in a specific health context. The unique nature of each person’s microbiota further complicates the matter, making it difficult to define a “normal” gut microbiota [40]. Furthermore, the lack of standardisation in testing methodologies results in inconsistent outcomes [44,45], limiting their utility in clinical practice. While microbiome tests hold promises for diagnosing specific conditions, their application in general healthcare remains limited, as they often provide little actionable information for diagnosis or treatment requiring highly specialised and microbiota-trained health professionals.

Among the potentially microbiota-modulating approaches that can be taken, the most well-known by the participants was faecal transplant (39.02% of the answers) [46] and the use pre- and probiotics (26.57% of the answers) [47]. This demonstrates an awareness of these options and opens opportunities to further educate on the advantages and disadvantages of different modulation strategies as the supporting science continues to develop.

In summary, this survey collected data on gut microbiota-related topics and the sources through which the general population and healthcare professionals are exposed to this information. It was clear that most respondents had received knowledge relating to the gut microbiota but had some misconceptions or incomplete knowledge, highlighting areas that could be addressed through science communication and dissemination. These include emphasising the multi-microorganism composition of the gut microbiota, factors that influence their composition, the various conditions associated with it, and more accurate and realistic information on gut microbiota tests and current modulation strategies.

Effective communication relating to the gut microbiota can educate healthcare professionals and broaden the repertoire of tools at their disposal to tackle patients’ conditions form a more holistic perspective, foster networking between scientists and the health system and educate the general public, while empowering the population to take control of their own health. To achieve this, citizen science initiatives [48,49] and interdisciplinary approaches between the science community, the health systems, public engagement agents and governmental administrations are needed to design effective strategies towards a more educated society [50].

## Acknowledgements

The authors want to thank Dr. Chloe Hutchins, Dr. Raul Cabrera Rubio and Kate O’Mahony for discussions; Macarena Forner, Beatriz Vera, Xavier García, Dr. Guilherme Martin, Tianqi Li, Prof. Leilei Yu, Dr. Ana Soriano, Dr. Coral Barcenilla, Dr. Laura Marroquí, Dr. Reinaldo Sousa Dos Santos, Dr. Natalia Gómez, Prof Paula Vitaglione, John Gallager, APC Microbiome Ireland and Teagasc for circulating the survey; Dr. Victor Strîmbu for his help with data analysis; and to all the participants for filling the survey.

## Author contributions

EG-G conceived, designed, conducted the study, wrote and edited the manuscript. SA designed, analysed data, wrote the manuscript and edited the manuscript. CO, SK, SD, BJ, VP, FHS, AM conducted the study. PDC conceived and edited the manuscript and secured funding. All authors contributed to the article and approved the submitted version.

## Statements and Declarations

### Ethical considerations

Not applicable as no personal data was collected.

### Consent to participate

Not applicable as no personal data was collected.

### Consent for publication

Not applicable.

### Data availability

The data set for all the answers in the different languages and the R computer codes used are available upon request.

