## Supplementary information for "Gut MicrobiotAware: how much do we know about gut microbiota? An international questionnaire"

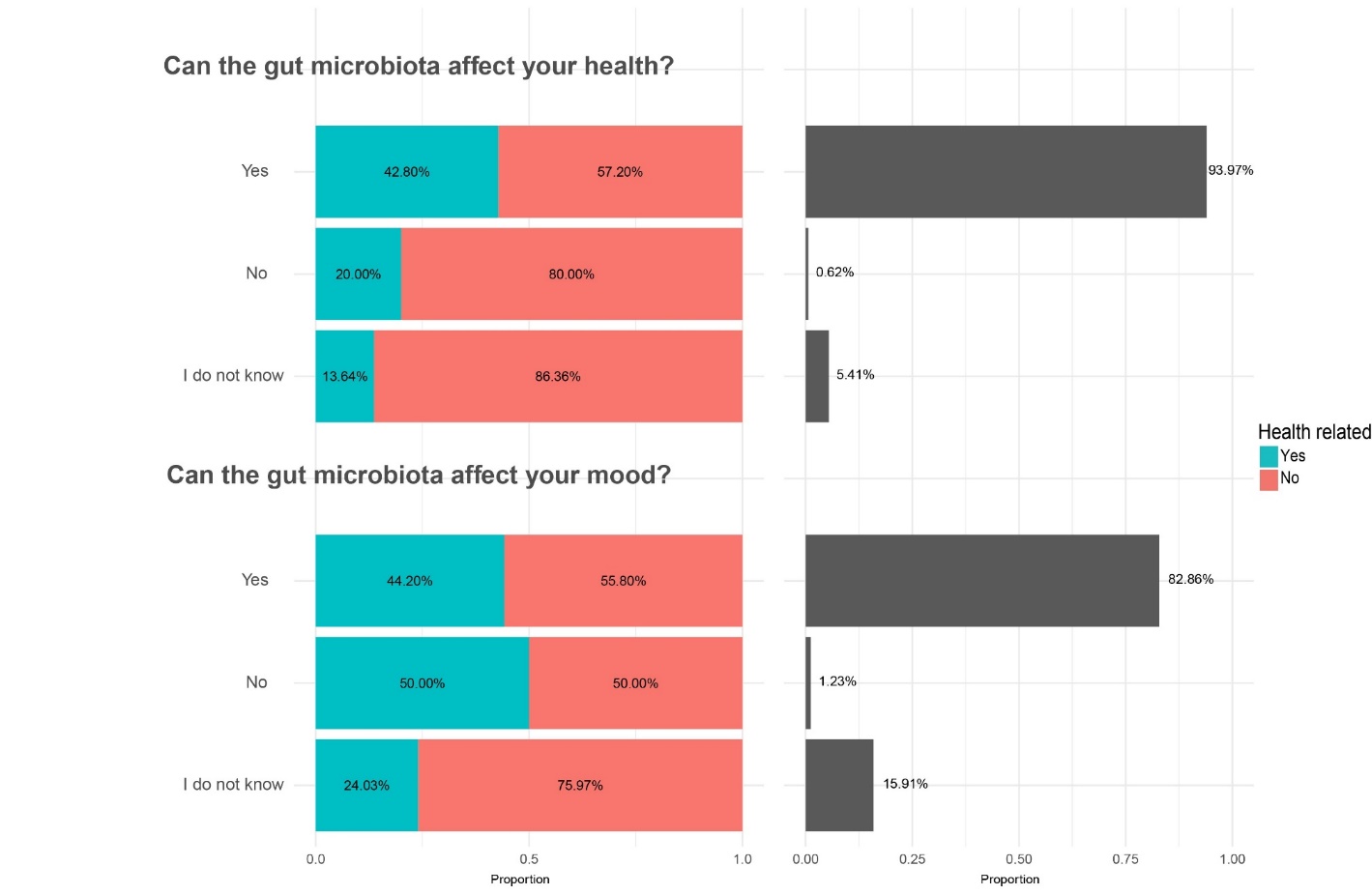
**Supplementary figures**

**Figure S1.** Participants' perceptions of whether the gut microbiota affects health and mood (n = 813, n = 811).

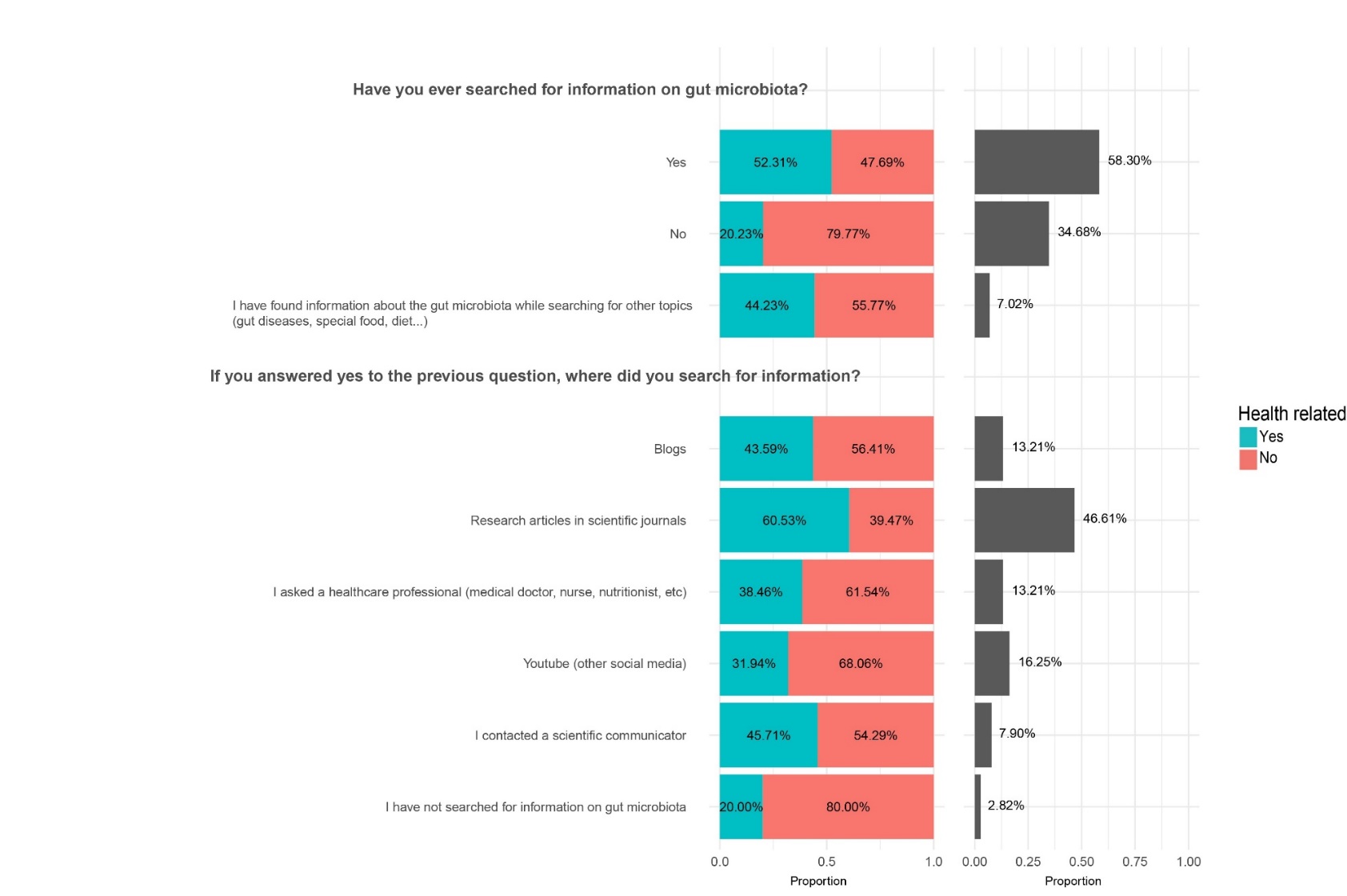

**Figure S2.** Information channels on gut microbiota that the participants had previously used (n = 741, n = 556).
